# Allele-specific open chromatin in human iPSC neurons elucidates functional non-coding disease variants

**DOI:** 10.1101/827048

**Authors:** Siwei Zhang, Hanwen Zhang, Min Qiao, Yifan Zhou, Siming Zhao, Alena Kozlova, Jianxin Shi, Alan R. Sanders, Gao Wang, Subhajit Sengupta, Siobhan West, Michael Streit, Chad A. Cowan, Mengjie Chen, Zhiping P. Pang, Pablo V. Gejman, Xin He, Jubao Duan

## Abstract

Functional interpretation of noncoding disease variants, which likely regulate gene expression, has been challenging. Chromatin accessibility strongly influences gene expression during neurodevelopment; however, to what extent genetic variants can alter chromatin accessibility in the context of brain disorders/traits is unknown. Using human induced pluripotent stem cell (iPSC)-derived neurons as a neurodevelopmental model, we identified abundant open-chromatin regions absent in adult brain samples and thousands of genetic variants exhibiting allele-specific open-chromatin (ASoC). ASoC variants are overrepresented in brain enhancers, transcription-factor-binding sites, and quantitative-trait-loci associated with gene expression, histone modification, and DNA methylation. Notably, compared to open chromatin regions and other commonly used functional annotations, neuronal ASoC variants showed much stronger enrichments of risk variants for various brain disorders/traits. Our study provides the first snapshot of the neuronal ASoC landscape and a powerful framework for prioritizing functional disease variants.

**One Sentence Summary:** Allele-specific open chromatin informs functional disease variants

## Main text

Most common disease risk variants implicated in genome-wide association studies (GWAS) are located in noncoding regions of the genome. Functional interpretation of putative causal variants at these loci is challenging. For neuropsychiatric disorders, despite recent progress on transcriptomic and epigenomic studies of human postmortem brains (e.g., PsychENCODE) (*1–4*), most disease causal variants/genes remain unknown. Because open/accessible chromatin often overlaps with regulatory DNA sequence (*5, 6*), co-localization of disease risk variants within open chromatin regions (OCRs) of human postmortem brains or iPSC-derived neurons can help prioritize putative functional noncoding risk variants for neuropsychiatric disorders (*7–10*). However, not all the variants within OCRs are functional, and as a result, the enrichment of GWAS signals in OCRs is often modest (*8*). It thus remains a challenge to precisely identify causal variants for neuropsychiatric disorders.

One strategy to address this challenge is to identify functional variants that affect chromatin accessibility which influences gene expression. In this work, we focus on allele-specific open chromatin (ASoC) variants that display allelic imbalance in sequencing reads at heterozygous single nucleotide polymorphism (SNP) sites. By comparing the chromatin accessibility of both alleles within the same sample, our approach minimizes experimental variations to better detect functional variants. Compared to another common approach of mapping functional variants, expression quantitative trait loci (eQTL), ASoC mapping has the advantage of directly identifying putatively functional variants, rather than those in linkage disequilibrium (LD). Despite these advantages, the landscape of ASoC in major neuronal cell types and its functional relevance to neuropsychiatric disorders such as schizophrenia (SZ), remain unknown.

Neuronal cells derived from iPSCs are a promising cellular model for neuropsychiatric disorders (*11, 12*), offering an alternative to human postmortem brains. iPSC can be derived from patients or healthy controls and then differentiated into different subtypes of neuronal cells of high purity in a controlled manner (*12, 13*). iPSC modelling may unravel unique functional genomic features pertaining to the developmental aspects of neuropsychiatric disorders, which may not be captured using postmortem brains. Here, we conducted a comprehensive mapping of ASoC variants in major iPSC-derived neuronal (iN) subtypes. These ASoC profiles provide a direct functional readout of noncoding risk variants of neuropsychiatric disorders.

We derived iPSC lines from 20 subjects selected from the Molecular Genetics of Schizophrenia (MGS) cohort (Table S1). These 20 subjects were chosen for being enriched for heterozygous (*i.e.*, informative for the allele-specific assay) index SNPs (*p* < 5 × 10^−8^) at ~70/108 SZ GWAS loci (*14, 15*) (Fig. S1, Methods). The iPSC lines were first differentiated into neural progenitor cells (NPC), and subsequently to early-stage (day-15) glutamatergic (iN-Glut) (*16*), GABAergic (iN-GA), and dopaminergic (iN-DN) (*17, 18*) neurons, with high purity (75-90%) (Fig. 1A, Fig. S2A-C). We carried out Assay for Transposase-Accessible Chromatin using sequencing (ATAC-seq) and RNA-sequencing (RNA-seq) in each cell type for 8 lines (“core 8” lines). To maximize the power of detecting ASoC for the study of disease GWAS variants, we performed ATAC-seq for 12 additional lines in NPCs and iN-Glut (Fig. 1A). We obtained 49-100M 50-bp paired-end (PE) ATAC-seq reads and 20-30 million 150-bp PE RNA-seq reads for each sample (Table S2). All analyzed ATAC-seq samples passed standard quality control based on the characteristic nucleosomal periodicity of fragment size distribution and high signal-to-noise ratio around transcription start sites (TSS) (Fig. S3). All ATAC-seq and RNA-seq samples were confirmed for individual identity using VerifyBamID, and the absence of chromosomal abnormality of each cell culture was verified by RNA-seq-based eSNP-karyotyping (Fig. S2D, Methods).

**Fig. 1.**
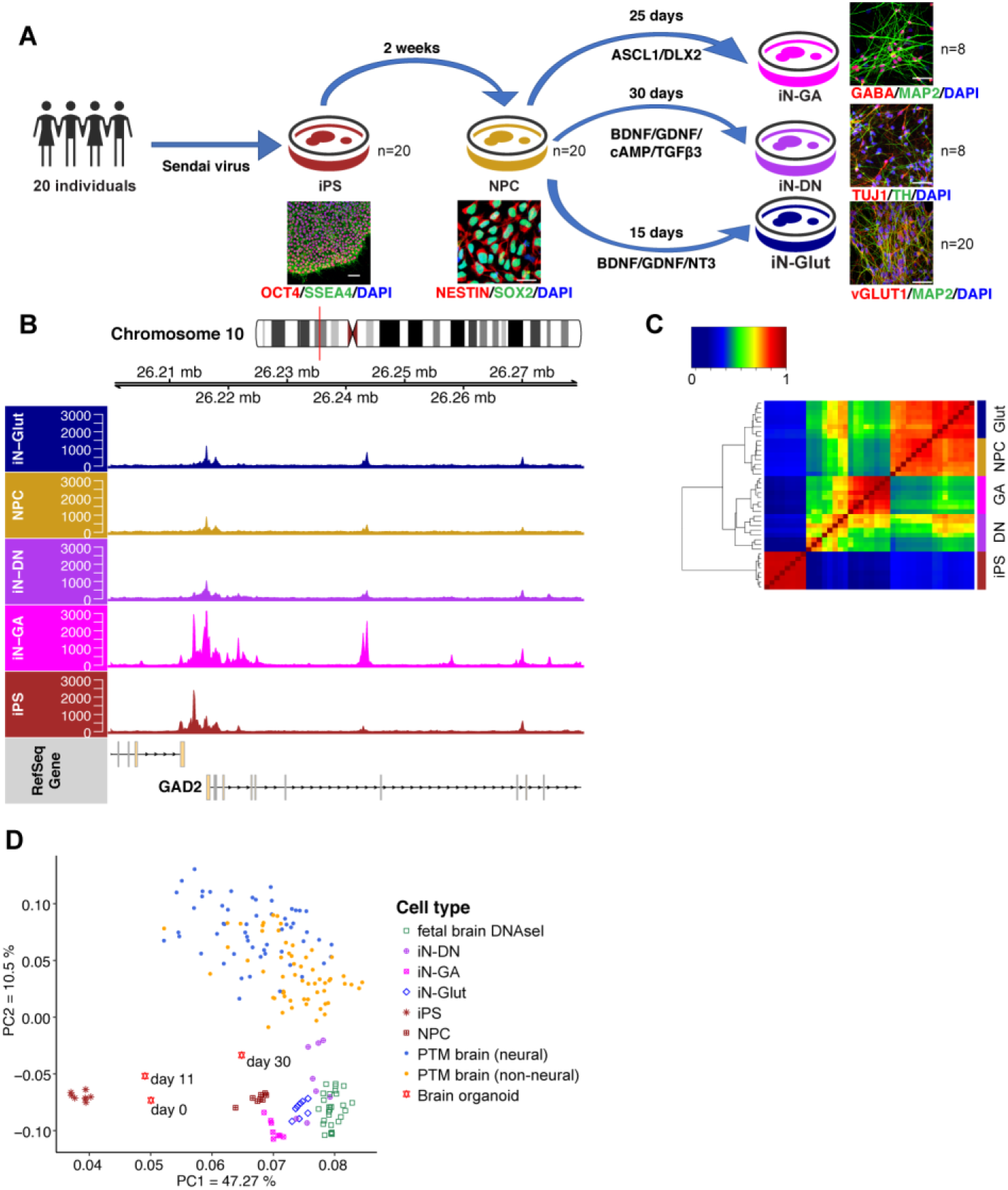
Mapping open chromatin regions (OCRs) by ATAC-seq in iPSC and neuronal cells. **(A)** Schematic of iPSC differentiation strategies. Scale bar: 100 μm in iPSC and 25 μm for others. **(B)** OCR peaks for *GAD2* in different cell types, showing increased chromatin accessibility in GABAergic neurons (iN-GA). **(C)** Hierarchical clustering and heatmap of normalized ATAC-seq reads within OCR peaks of different cell types. **(D)** Principal component analysis (PCA) of OCR peak intensities shows the separation of different cell types/tissues as well as the higher similarity between iNs and fetal brains/day-30 fetal organoids than postmortem (PTM) adult brain. All data were ATAC-seq except for fetal brain being DNase-seq.

To call OCR peaks, we pooled the reads of all 8 ATAC-seq samples within each cell type. We identified 256~337 K OCR peaks (FDR< 0.05) in each cell type (Fig. S4A, Table S3). iPSCs had the highest number of peaks, consistent with the highly permissive chromatin state in pluripotent stem cells (*19*). Peak intensities (normalized ATAC-seq read counts) were highly correlated between samples within a cell type (R > 0.9) (Fig. S4B), confirming the high data quality. We estimated that the total peaks called from 8 samples accounted for ~70% of all possible peaks (Fig. S4C, Methods). The median OCR length was 335 bp, and OCR peaks in each cell type covered 4-5% of the genome (Fig. S4D, Table S3).

We first confirmed the OCRs for some known cell-type-specific genes, such as *GAD2*, encoding an enzyme responsible for the synthesis of GABA in iN-GA, and *NANOG*, encoding a transcription factor (TF) essential for maintaining pluripotency in iPSCs (Figs. 1B and S4E). We then clustered 40 ATAC-seq samples (8 samples each × 5 cell types) using quantile-normalized reads of a common set of 666,614 non-overlapping peaks. We found strong cell-type-specific clustering, consistent with that from using RNA-seq data (Figs. 1C and S5A-C). Using the same set of OCR peak intervals, we further carried out principal component analysis (PCA) to compare our ATAC-seq data with publicly available open chromatin datasets from fetal cortical organoids (*3*), fetal brains (*20*), and PsychENCODE adult brains (*8*). We found that iPSCs formed a most distant cluster, while our iPSC-derived neural cells were more similar to fetal brains and day-30 cortical organoids than to adult brains (Fig. 1D), suggesting our dataset better captures early stages of neurodevelopment than adult brains.

We next evaluated whether our neuronal OCR peaks were comparable to PsychENCODE brain ATAC-seq peaks (*n* = 117,935) (*8*). We defined the overlapping peaks as those reciprocally sharing at least 25% of their peak intervals. We found that OCRs of individual neuronal cell types (NPC, Glut, DN, or GA) overlapped with 45-55% of the PsychENCODE brain peaks (64% for the combined neuronal peaks, Table S3). However, peaks overlapping with PsychENCODE data only account for ~20% of our OCRs, *i.e.*, most of our neuronal OCRs are not found in PsychENCODE brains. Using a more stringent peak calling cut-off (FDR < 1%) gave similar results (Table S3). Our result is consistent with fetal cortex possessing about two-fold more enhancers than developed cerebral cortex (*3*). These observations suggest that OCRs in our iPSC cellular model capture a majority of regulatory elements in adult brain, and also expand the repertoire of regulatory elements activated only during early development.

To identify SNPs that may affect chromatin accessibility and possibly regulate gene expression, we tested which heterozygous SNPs exhibited allelic imbalance of ATAC-seq reads (*i.e.*, ASoC) in each cell type (Fig. S6, Methods). We observed that the directionality of the allelic imbalance of candidate ASoC SNPs (binomial *p* < 0.05) was highly concordant between individuals (Fig. S7A), and thus we adopted a pooling approach (*21, 22*) to increase the power to detect ASoC. Specifically, we pooled the allele counts of heterozygous SNPs of the core 8 cell lines for each cell type to call ASoC (Fig. S6). We identified 920~2,392 ASoC SNPs in each cell type (Fig. 2A, Tables S4-8). Most OCR peaks contained a single ASoC SNP (Fig. S7B, Table S9). The proportion of heterozygous SNPs showing ASoC was similar in each cell type (~1.5%) except for iN-Glut (3.9%), and the larger number of ASoC SNPs in iN-Glut (*n* = 2,392) was unlikely due to the difference of sequencing depth (Fig. S7C, Table S11).

**Fig. 2.**
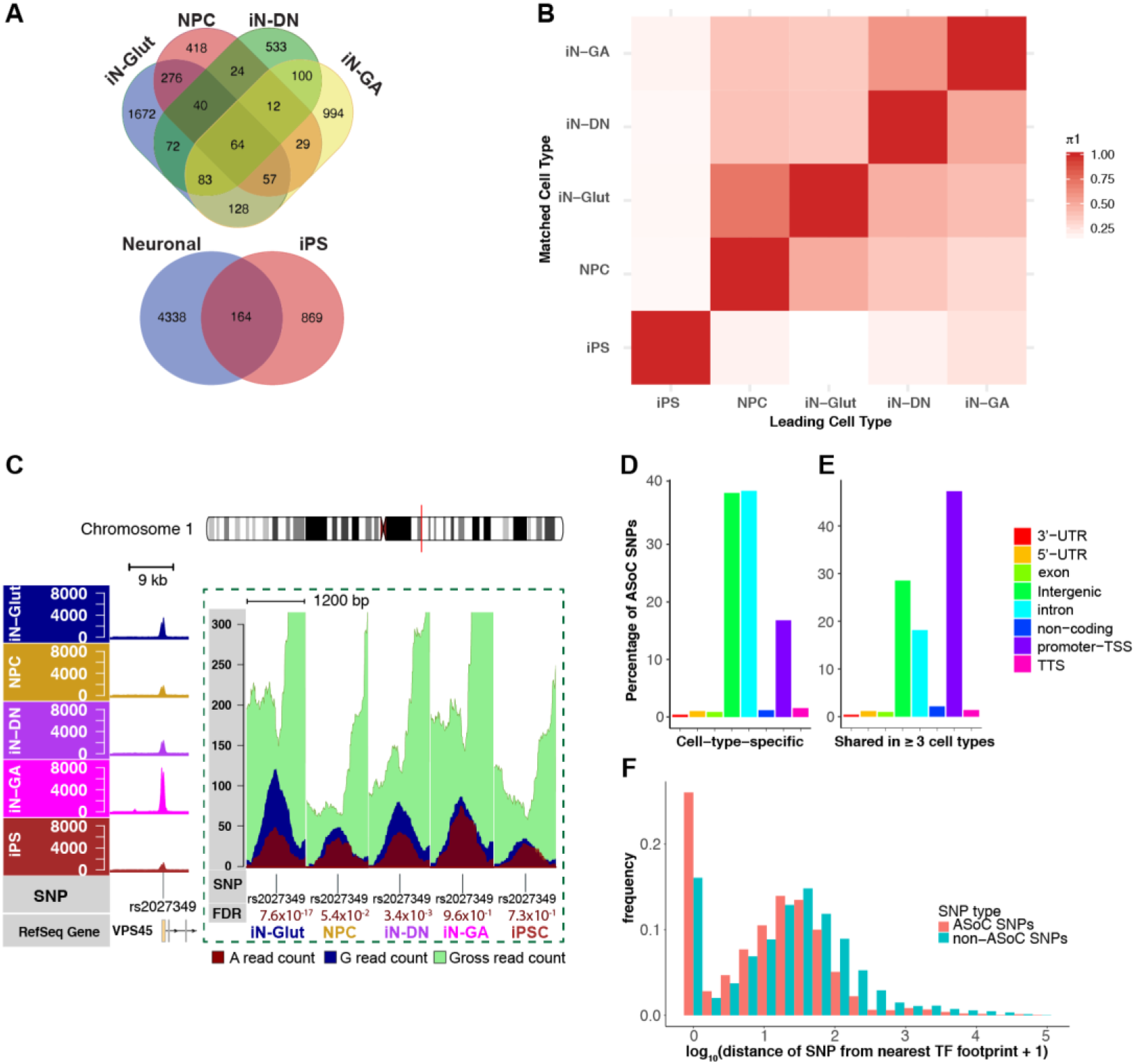
Characteristics of ASoC SNPs. **(A)** Venn diagram of ASoC SNP numbers (*n* = 8 samples per cell type). **(B)** Heatmap of pairwise π1 estimation of ASoC SNPs among five cell types. Each number in the matrix represents the proportion of non-null tests (π1) estimated from binomial *p* values in the matched cell type for ASoC SNPs (FDR < 0.05) obtained from the leading cell type. **(C)** ATAC-seq peaks near rs2027349, the strongest neuronal ASoC SNP that is also associated with SZ, in each cell type. The high-resolution read pileups flanking the SNP (from 4 heterozygous samples) and allelic ratio differences are shown in the dashed box. **(D-E)** The proportion of ASoC SNPs that are cell type-specific (D) or shared between at least three cell types (E) in different types of genomic regions. **(F)** Histogram of the ASoC SNPs based on their distances (bp) to the nearest TF footprint.

The ASoC data allowed us to investigate how regulatory variants differ across 5 cell types. Simple comparison of ASoC SNPs (at FDR < 0.05) between cell types revealed abundant cell-type-specific ASoC SNPs, in particular between neuronal cell types and iPSC (Fig. 2A). However, this analysis may underestimate the extent of ASoC sharing, because the detection power of ASoC in each cell type is less than 100%. Similar to cross-tissue eQTL studies (*23, 24*), we used Storey’s π1 analysis to estimate pairwise ASoC sharing (*25*). We ascertained ASoC SNPs in one cell type (FDR < 0.05) and estimated the fraction of them that are also ASoC in the second cell type (Fig. 2B). As expected, we found that neuronal cell types shared a much higher percentage of ASoC SNPs with each other than with iPSC (30~70% vs. 10~20%, Fig. 2B). Even among neuronal cells, there were large fractions of ASoC SNPs unique to one cell type. For SNPs that were identified as ASoC in multiple cell types, the direction of allelic imbalance was highly correlated between cell types (R = 0.77~0.95; Fig. S7D), suggesting conserved regulatory mechanisms of chromatin accessibility for a large subset of ASoC SNPs across cell types.

The observed abundance of cell-type specific ASoC SNPs may be driven by either cell-type-specific OCRs or different SNP effect sizes (*i.e.*, allelic ratios) across cell types. To distinguish these two possible mechanisms, we compared ASoCs in each neuronal cell type vs. iPSCs. We ascertained neuron-specific ASoC SNPs as those with FDR < 5% in one neuronal cell type but had low read depth (<20 reads) or insignificant allele imbalance (binomial test *p* > 0.05) in iPSCs (Fig. S8). We found that about 38~52% of neuron-specific ASoCs were associated with OCRs that are likely neuron-specific, defined as > 2-fold (at FDR < 5%) higher peak intensity than that in iPSC (Fig. S9). On the other hand, a large number of neuron-specific ASoC SNPs also showed insignificant allele imbalance in iPSCs, despite high sequencing depth (> 100 reads) in iPSC (Fig. S8), reflecting cell-type-specific allelic effect size. For example, the neuron-specific ASoC SNP that showed the strongest association with SZ, rs2027349 in the 5’-UTR of *VPS45*, exhibited ASoC in iN-Glut, iN-DN, and NPC but not in iPSCs or iN-GA, largely due to different allelic effect sizes across cell types (Fig. 2C). These results suggest that variations of both OCR intensities and SNP effect sizes play play significant roles in driving cell-type specific ASoC.

We next compared the genomic/epigenomic features of shared vs. cell-type specific ASoC variants. For this analysis, we defined shared ASoC variants as those found in three or more cell types, and cell-type specific variants as those unique to one cell type. While ~80% of cell-type-specific ASoC SNPs were found to be intergenic or intronic, the majority of the shared ASoC SNPs were in promoter-TSS regions (Fig. 2D-E). We further annotated SNPs using ChromHMM-based genomic features in brain such as promoters and enhancers (*7*). We found a higher percentage of cell-type-specific ASoC SNPs (30% vs. 16% of the shared SNPs) in enhancers (Fig. S10A). Using a binomial test implemented in GREAT (*26*), we found that both types of ASoC SNPs were enriched in promoters and enhancers (Fig. S10B). This was not unexpected, because differentially accessible chromatin regions across brain regions were also found to be enriched for both promoters and enhancers (*27*). Our analysis of differentially accessible OCRs across cell types revealed similar finding (Fig. S4F-H). However, compared to shared ASoCs, cell-type-specific ASoCs showed ~2-fold higher enrichment in enhancers overall (Fig. S10B), and an overall stronger enrichment in different subtypes of brain enhancers (Fig. S10C-F). Furthermore, comparing the relative distribution of ASoC SNPs in enhancers vs. promoters (Fig. S10A), cell-type-specific ASoC SNPs were found strongly enriched in enhancers (odds ratio [OR] = 2.13, *p*=7.3 × 10^−7^, Fisher’s Exact Test), while the shared ASoCs were enriched in promoters (OR = 4.6, *p*=4.3 × 10^−35^, Fisher’s Exact Test). These results suggest that ASoC SNPs are often associated with regulatory activities in the brain, with cell-type-specific ASoCs more associated with enhancer activities.

A major mechanism of regulatory variants is alternation of chromatin accessibility to TFs (*5, 28*). We thus mapped TF-binding site (TFBS) footprints from ATAC-seq data and examined their enrichment of ASoC SNPs (Methods). In iN-Glut neurons, we identified 612,373 TFBSs for a set of 579 TF motifs from the JASPAR Vertebrate database (*29*). Out of the 2,392 ASoC SNPs, 622 (26%) were found inside TFBSs, representing a 1.8-fold enrichment (vs. non-ASoC SNPs, *p* = 9.1×10^−34^, Fisher’s exact test; Fig. 2F). Other cell types also gave similar results (Fig. S11). This is consistent with a previous report that only 12% of the genetic variants affecting TF-binding are located in TF-binding motifs (*30*). However, most ASoC variants are still within 200 bp of the nearest footprint (Fig. 2F), an interval enriched with SNPs showing allele-specific TF occupancy (*30*), suggesting ASoC variants may commonly affect TF-binding; although other mechanisms may also play roles.

To explore the relevance of neuronal ASoC SNPs to gene regulation in the brain, we jointly analyzed our data with QTL data for other molecular phenotypes in the brain. As expected, the number of detected ASoC SNPs linearly increases with sample size (Fig. S12A), we thus used ASoC SNPs of all 20 lines (for NPC and iN-Glut) to maximize the power. We identified 5,611 and 3,547 ASoC variants (1,690 shared, FDR < 5%) in iN-Glut and NPCs, respectively (Fig. 3A, Tables S12-13). We first explored the regulatory relevance of these ASoC variants using PsychENCODE high-confidence enhancers of fetal brain organoids and adult brains (*1, 3*). We found ASoC SNPs in both iN-Glut and NPCs were enriched in fetal brain organoid enhancers (*p* = 1.8×10^−12^ and 2.4×10^−24^ respectively; Fisher’s exact test). Interestingly, ASoC SNPs in iN-Glut, but not NPCs, showed enrichment in adult brain enhancers (*p* = 9.5×10^−9^, Fisher’s exact test, Fig. S12B, Tables S12-14), consistent with the fact that iN-Glut represents a more mature neuronal state than NPCs. We next assessed the enrichment of ASoC SNPs for regulatory variants associated with gene expression, histone modification, and DNA methylation from an adult brain QTL study (Fig. 3B, Tables S15-16) (*31*). We focused on iN-Glut because of its relevance to SZ (*32, 33*) and its stronger enrichment (vs. NPC) in adult brain enhancers (Fig. S12B). Enrichment analysis of QTL data, however, faces two challenges: SNPs passing statistical cut-offs may not be causal variants but their LD proxies, and many causal SNPs do not pass stringent statistical cutoffs. To address these issues, we used TORUS (*34*), a tool based on a Bayesian hierarchical model, to perform the enrichment analysis, as recently done in large eQTL studies (*24*). We found that ASoC SNPs were highly enriched (30~90-fold) for putatively causal QTL variants of expression (eQTL), histone marks (haQTL), and DNA methylation (meQTL) (Fig. 3B). The observed enrichments of iN-Glut ASoC SNPs for brain QTLs were also confirmed by using independent eQTL and meQTL data sets (Fig. S12D) (*1, 8, 35*). Altogether, our results support the regulatory effects of ASoC SNPs and suggest that some neuronal ASoC variants may have lasting functions at later developmental stages.

**Fig. 3.**
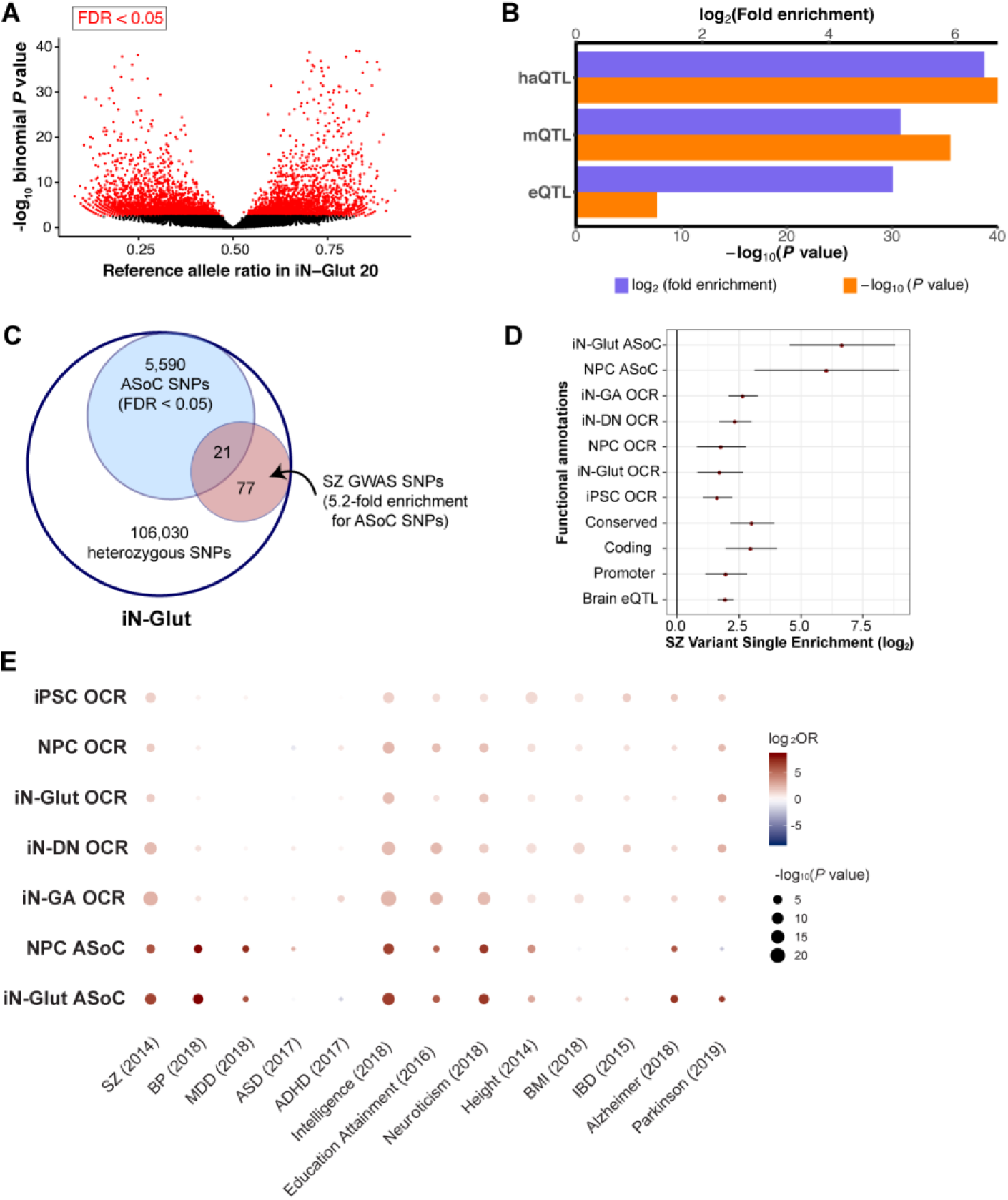
Enrichments of ASoC variants for brain QTLs and genetic risk of SZ and other complex disorders/traits. **(A)** Volcano plot of the reference allelic ratio of all heterozygous SNPs from 20 iN-Glut samples. Red: ASoC SNPs (FDR<5%). **(B)** Enrichment of iN-Glut ASoC SNPs for brain QTLs (eQTL: expression QTL, mQTL: DNA methylation QTL, haQTL: histone acetylation QTL). **(C)** Venn diagram of all heterozygous SNPs and ASoC SNPs in 20 iN-Glut samples, showing the enrichment ASoC SNPs for SZ GWAS SNPs. **(D)** Enrichments of for SZ GWAS SNPs for different annotations after accounting for SNP LD. The red dot indicates the log2 odds ratio of enrichment, and the bar-range represents 95% confidence interval. **(E)** Bubble plot showing the enrichment of GWAS associations of 13 disorders/traits (see Methods) in ASoC SNPs and OCRs. Dot color indicates the log2 odds ratio of enrichment, and dot size denotes its corresponding significance. SZ: Schizophrenia, BP: Bipolar, MDD: Major depression disorder, ADHD: Attention-deficit/hyperactivity Disorder, BMI: Body mass index, IBD: Inflammatory bowel disease.

Having shown the functional relevance of neuronal ASoC SNPs in the brain, we assessed the utility of ASoC in inferring functional noncoding risk variants for neuropsychiatric disorders. We and others have previously shown that disease risk variants are modestly over-represented in OCRs (*10, 36, 37*). We hypothesized here that ASoC variants were further enriched for functional disease variants. We first examined the enrichment of iN-Glut ASoC SNPs (*n* = 20 lines) in variants associated with SZ (*15*) (Fig. 3C, Tables S17-18). We found 21 of the 5,611 ASoC SNPs were GWAS index SNPs or their LD proxies at 17 independent SZ loci (nearest genes: *VPS45, BCL11B, GALNT10, RERE, UBE2Q2P1, PCDHA1, NGEF, DPYD*, *BAG5, BTA3A2, LOC100507431, KMT2E-AS1, PBRM1, ZSCAN16-AS1, STAT6, LINC000637* and *PPP1R16B)*, representing a 5.2-fold enrichment (*p* = 1.5×10^−8^, Fisher’s exact test, Fig. 3C). To account for LD and uncertainty of causal variants as described above for the QTL enrichment analysis, we applied TORUS (*34*) to SZ GWAS summary statistics (*15*) (Fig. 3D, Methods). We found that neuronal (iN-Glut or NPC) ASoC SNPs showed remarkably higher enrichment for SZ risk variants than OCRs and other functional annotations, such as conservation and brain eQTL (50~90 vs. 2.5~6-fold, Fig. 3D).

We expanded the TORUS enrichment analysis of GWAS risk variants to 9 other brain disorders and traits (Fig. 3E, Fig. S13). Similar to SZ, we observed much higher enrichment of GWAS variants for intelligence, educational attainment, and neuroticism in both iN-Glut and NPC ASoC variants than in OCRs. For bipolar disorder (BP) and major depressive disorder (MDD), we observed strong enrichment of their respective GWAS variants in neuronal ASoC SNPs, but not in OCRs. For neurodegenerative disorders, we found significant enrichment of ASoC SNPs (but only in iN-Glut) in Alzheimer’s disease (AD), but not Parkinson’s disease (PD), although OCRs did show weak enrichment for PD. For comparison, we included GWAS datasets of body mass index (BMI), height, and inflammatory bowel disease (IBD). BMI and IBD variants showed no or low enrichment in neuronal ASoC SNPs or OCRs. We observed strong enrichment of height risk variants in neuronal ASoC SNPs, which may reflect the highly polygenic nature of height and the genetic correlation between human height and intelligence (*38*). We note that the failure to detect enrichment in ASoC or OCRs for some diseases may reflect underpowered disease GWAS (*e.g.*, attention deficit hyperactivity disorder [ADHD] and autism spectrum disorder [ASD]), weaker disease relevance of the assayed cell types (*e.g.*, for AD and PD), or the limited power of our study with only 20 cell lines. Nonetheless, neuronal ASoC SNPs overall showed much stronger enrichment of risk variants for brain-related phenotypes than other commonly used annotations such as OCRs and conservation (Figs. S12C and S13). Together, these results suggest that ASoC is a highly effective predictor of functional noncoding variants for various brain disorders and traits.

In summary, we have provided the first snapshot of the ASoC landscape in an iPSC-based neurodevelopmental model and demonstrated that ASoC is a direct functional readout of noncoding risk variants of brain disorders/traits. The enrichments of neuronal ASoC SNPs for brain enhancers, TFBSs, and brain QTLs suggest mechanistic links between chromatin accessibility and gene expression. Given the strong enrichment of ASoC variants for GWAS signals of SZ and other brain disorders/traits, our study provides a useful resource and perhaps more importantly, an effective framework for functional interpretation of noncoding disease risk variants.

## Acknowledgements

We thank Dr. William J. Greenleaf at Stanford for his help on ATAC-seq analyses. This work was supported by NIH grants: R01MH106575, R01MH116281, and R01AG063175 (JD); R01MH110531 (XH).

## Authors contributions

S.Z. analyzed the ATAC-seq, bulk, and scRNA-seq data, and wrote the manuscript. H.Z. performed the experiments, analyzed data, and wrote the manuscript. M.Q., S.Z., Y.Z., and G.W., performed computational analyses of tissue-specific ASoC, TF footprints, GWAS enrichment, fine-mapping and CROP-seq. A.K. and M.S. carried out DA neuron differentiation. J.S. and S.S. helped with data QC and statistical analyses. S.W. helped with CROP-seq gRNA design and library preparation. A.R.S. and P.V.G. helped with clinical phenotypes, data interpretation, and manuscript writing. C.A.C. and Z.P.P. guided the gene editing and neuron differentiation, respectively. M.C. helped with the CROP-seq data analysis. X.H. supervised the data analyses of M.Q., S.Z., and Y.Z., and wrote the manuscript. J.D. conceived the study, supervised the project, and wrote the manuscript.

## References

1. M. J. Gandal et al., Transcriptome-wide isoform-level dysregulation in ASD, schizophrenia, and bipolar disorder. Science 362, (2018).

2. P. Rajarajan et al., Neuron-specific signatures in the chromosomal connectome associated with schizophrenia risk. Science 362, (2018).

3. A. Amiri et al., Transcriptome and epigenome landscape of human cortical development modeled in organoids. Science 362, (2018).

4. M. Li et al., Integrative functional genomic analysis of human brain development and neuropsychiatric risks. Science 362, (2018).

5. J. F. Degner et al., DNase I sensitivity QTLs are a major determinant of human expression variation. Nature 482, 390–394 (2012).

6. R. E. Thurman et al., The accessible chromatin landscape of the human genome. Nature 489, 75–82 (2012).

7. L. de la Torre-Ubieta et al., The Dynamic Landscape of Open Chromatin during Human Cortical Neurogenesis. Cell 172, 289–304 e218 (2018).

8. J. Bryois et al., Evaluation of chromatin accessibility in prefrontal cortex of individuals with schizophrenia. Nat Commun 9, 3121 (2018).

9. J. F. Fullard et al., An atlas of chromatin accessibility in the adult human brain. Genome research 28, 1243–1252 (2018).

10. M. P. Forrest et al., Open Chromatin Profiling in hiPSC-Derived Neurons Prioritizes Functional Noncoding Psychiatric Risk Variants and Highlights Neurodevelopmental Loci. Cell Stem Cell 21, 305–318 e308 (2017).

11. D. M. Panchision, Concise Review: Progress and Challenges in Using Human Stem Cells for Biological and Therapeutics Discovery: Neuropsychiatric Disorders. Stem Cells 34, 523–536 (2016).

12. Z. Wen, K. M. Christian, H. Song, G. L. Ming, Modeling psychiatric disorders with patient-derived iPSCs. Current opinion in neurobiology 36, 118–127 (2016).

13. K. J. Brennand et al., Modelling schizophrenia using human induced pluripotent stem cells. Nature 473, 221–225 (2011).

14. J. Shi et al., Common variants on chromosome 6p22.1 are associated with schizophrenia. Nature 460, 753–757 (2009).

15. C. Schizophrenia Working Group of the Psychiatric Genomics, Biological insights from 108 schizophrenia-associated genetic loci. Nature 511, 421–427 (2014).

16. Z. Wen et al., Synaptic dysregulation in a human iPS cell model of mental disorders. Nature 515, 414–418 (2014).

17. R. Gonzalez et al., Deriving dopaminergic neurons for clinical use. A practical approach. Sci Rep 3, 1463 (2013).

18. S. Kriks et al., Dopamine neurons derived from human ES cells efficiently engraft in animal models of Parkinson’s disease. Nature 480, 547–551 (2011).

19. E. Meshorer, T. Misteli, Chromatin in pluripotent embryonic stem cells and differentiation. Nat Rev Mol Cell Biol 7, 540–546 (2006).

20. B. E. Bernstein et al., The NIH Roadmap Epigenomics Mapping Consortium. Nat Biotechnol 28, 1045–1048 (2010).

21. V. Onuchic et al., Allele-specific epigenome maps reveal sequence-dependent stochastic switching at regulatory loci. Science 361, (2018).

22. J. Xu et al., Landscape of monoallelic DNA accessibility in mouse embryonic stem cells and neural progenitor cells. Nat Genet 49, 377–386 (2017).

23. E. Grundberg et al., Mapping cis- and trans-regulatory effects across multiple tissues in twins. Nat Genet 44, 1084–1089 (2012).

24. G. T. Consortium et al., Genetic effects on gene expression across human tissues. Nature 550, 204–213 (2017).

25. J. D. Storey, R. Tibshirani, Statistical significance for genomewide studies. Proc Natl Acad Sci U S A 100, 9440–9445 (2003).

26. C. Y. McLean et al., GREAT improves functional interpretation of cis-regulatory regions. Nat Biotechnol 28, 495–501 (2010).

27. R. E. Handsaker et al., Large multiallelic copy number variations in humans. Nat Genet, (2015).

28. M. R. Corces et al., The chromatin accessibility landscape of primary human cancers. Science 362, (2018).

29. A. Khan et al., JASPAR 2018: update of the open-access database of transcription factor binding profiles and its web framework. Nucleic Acids Res 46, D1284 (2018).

30. T. E. Reddy et al., Effects of sequence variation on differential allelic transcription factor occupancy and gene expression. Genome research 22, 860–869 (2012).

31. B. Ng et al., An xQTL map integrates the genetic architecture of the human brain’s transcriptome and epigenome. Nat Neurosci 20, 1418–1426 (2017).

32. D. Wang et al., Comprehensive functional genomic resource and integrative model for the human brain. Science 362, (2018).

33. H. K. Finucane et al., Heritability enrichment of specifically expressed genes identifies disease-relevant tissues and cell types. Nat Genet 50, 621–629 (2018).

34. X. Wen, Molecular QTL discovery incorporating genomic annotations using Bayesian false discovery rate control. Ann Appl Stat 10, (2016).

35. A. E. Jaffe et al., Mapping DNA methylation across development, genotype and schizophrenia in the human frontal cortex. Nat Neurosci 19, 40–47 (2016).

36. J. K. Pickrell, Joint analysis of functional genomic data and genome-wide association studies of 18 human traits. Am J Hum Genet 94, 559–573 (2014).

37. H. K. Finucane et al., Partitioning heritability by functional annotation using genome-wide association summary statistics. Nat Genet 47, 1228–1235 (2015).

38. M. C. Keller et al., The genetic correlation between height and IQ: shared genes or assortative mating? PLoS genetics 9, e1003451 (2013).

